# Genome-aware annotation of CRISPR guides validates targets in variant cell lines and enhances discovery in screens

**DOI:** 10.1101/2024.01.14.575203

**Authors:** Simon Lam, John C. Thomas, Stephen P. Jackson

## Abstract

Pooled CRISPR-Cas9 genetic knockout screens are powerful high-throughput tools for identifying chemo-genetic, synthetic-lethal and synthetic-viability interactions and are used as a key step towards identifying disease-modifying knockout candidates and informing drug design and therapeutic regimens. CRISPR guide libraries are commercially available for purchase and have been widely applied in different cell lines. However, discrepancies between the genomes used to design CRISPR libraries and the genomes of the cells subjected to CRISPR screens lead to loss of signal or introduction of bias towards the most conserved genes. Here, we present an algorithm, EXOme-guided Reannotation of nuCleotIde SEquences (Exorcise), which uses sequence search and CRISPR target annotation to adapt existing CRISPR libraries to user-defined genomes and exomes. Applying Exorcise on 55 commercially available CRISPR-spCas9 knockout libraries for human and mouse, we found that all libraries have mis-annotations, and that design strategy affects off-target effects and targeting accuracy relative to a standard reference sequence. In simulations on synthetic data, we modelled common mis-annotations in CRISPR libraries and found that they adversely affected recovery of the ground truth for all genes except for those with the strongest signals. Finally, we reanalysed DepMap and DDRcs CRISPR screens with Exorcise annotations and found that strong hits were retained, and lower-confidence hits were strengthened. Use of Exorcise on DepMap with exomes inferred from transcriptomic expression data demonstrated that cell-line–aware reannotation is possible without whole-genome sequencing. Taken together, our results show that Exorcise is a powerful reannotation tool that focuses existing CRISPR libraries towards the cell line genome under investigation and allows post-hoc reanalysis of completed CRISPR screens. Exorcise is open-source software licenced under a Creative Commons Zero Universal 1.0 licence and is available at <https://github.com/SimonLammmm/exorcise>.

## Introduction

Chemo-genetic relationships describe the modification of drug response as a function of genotype. Genetic knockout or gene silencing may confer resistance or hypersensitivity to a drug, and changes in expression of one gene can alter the effect of other genes on the phenotype. Technologies such as CRISPR-Cas9 and RNA interference can be used to interrogate gene-gene relationships and drug-gene interactions in a high-throughput manner using libraries of guide RNAs targeting precise genomic loci with sequence specificity and observing the drug modifying effect of CRISPR targeting. Drug-gene interactions are valuable and have potential medical applications for drug development, selective disease tissue targeting, studying cancer evolution of drug resistance, and for patient stratification and personalised medicine selection based on the variants present^1,2^.

The CRISPR-Cas9 system is a powerful tool to introduce precise genetic knockouts^3^. In CRISPR-Cas9, guide RNAs recruit the Cas9 endonuclease to target genomic loci through base-pairing of the guide with the cell’s genomic DNA and direct interaction between Cas9 and the protospacer-adjacent motif (PAM), which is 5’-NGG-3’ in the case of *Streptococcus pyogenes* Cas9 (spCas9), where N can be any of A, C, G, and T. Recruitment of spCas9 to DNA results in a precise double-strand break between the third and fourth nucleotide upstream from the PAM. These DNA breaks are then frequently repaired by error-prone nonconservative end-joining repair mechanisms, leading to small insertions and deletions. In specific genes, these can lead to frameshift nonsense mutations and functional knockout of the gene. In a pooled CRISPR-Cas9 screen, guides are introduced into cells as a pool, resulting in a population of cells with different genetic knockouts. Cells can then be treated with a chemical challenge. Cells with a knockout that confers resistance to the chemical have a fitness advantage and therefore proliferate at a higher rate than control; cells with a knockout that confers hypersensitivity to the chemical die out or proliferate slower than control. Guide abundance in the cell population during a screen can be measured by next-generation sequencing and compared between different timepoints or treatment conditions. In chemo-genetic screens, this provides a readout of relative guide abundances and definition of those whose action in knocking out their target gene conferred drug resistance, which would be enriched in the drug-treated population compared to control, and guides whose knockout conferred hypersensitivity, which would be depleted in the drug-treated population compared to control^4^.

CRISPR-Cas9 libraries contain guides with ∼20-nucleotide spacer sequences providing the base-pairing with the target region. In a random human-sized genome, this design provides sufficient precision to uniquely specify a target without off-target effects. However, repetitive sequences and gene duplications mean that off-target events occur more often than would be expected in a random genome. Other groups have previously studied CRISPR editing efficiency and off-target prediction relative to guide content, genome sequence context, and open-reading frame cutting consequence^5–7^, but the extent of off-target events in commercially available libraries and on-target specificity in non-standard genome assemblies, such as cancer cell lines, are yet to be inspected.

Here, we present Exorcise (EXOme-guided Reannotation of nuCleotIde SEquences), an algorithm that reannotates CRISPR screen guides by alignment to a user-defined genome and exon annotation on the same coordinate system (**Figure 1**). The program aligns guides to the genome and checks for Cas9 cut sites within exon boundaries. Exorcise detects the following events for each guide in a library: 1) off-target effects, where a guide targets exons within more than one gene; 2) missed-target effects, where a guide does not engage with its prescribed target; and 3) false non-targeting effects, where a valid guide-target annotation is missing. The algorithm does not depend on prior annotations and is able to annotate guides from arbitrary user-defined genomes and exomes alone. Using this tool, we found that 55 commercially available pooled CRISPR knockout screen libraries contain guides with missed-target effects in RefSeq genes. In a synthetic dataset, we simulated a CRISPR screen with a known ground truth and found that common types of mis-annotation found in commercially available libraries confer adverse effects on recovery of the ground truth but Exorcise ameliorates this by correcting the mis-annotations. Finally, we reanalysed DepMap CRISPR screens with Exorcise against RefSeq exons and exomes inferred from transcriptomes, separately, and found that the transcriptome-based reannotation is sufficient where whole-genome reconstruction of cancer cell lines is not possible. We therefore present Exorcise as a method to validate CRISPR targets given a supplied genome and exome, and to enhance discoveries in previously performed CRISPR screens without redesigning libraries or repeating experiments. Exorcise is available as an open-source computer program under a Creative Commons Zero 1.0 Universal licence.

**Figure 1.**
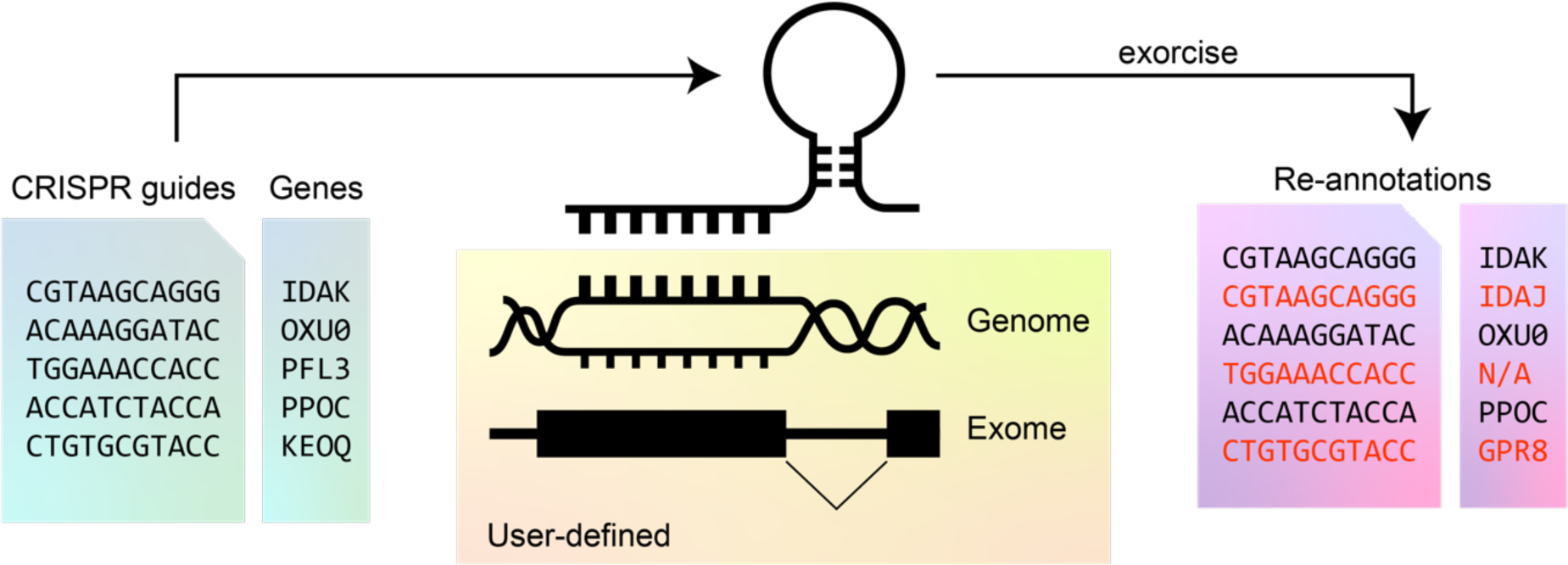
Overview of Exorcise. Library guides are aligned to a user-defined genome and annotated with a user-defined exome. Reannotations that differ from original annotations are shown in red.

## Results

### Off-target effects are common in CRISPR libraries

CRISPR targets are specified by base-pairing between guide RNAs and endogenous DNA sequences. Ideal guides specify their targets uniquely to ensure that effects on phenotype are causally related to CRISPR knockout at a single genomic locus. Off-target effects occur when guides result in CRISPR knockout of an unintended gene. We used Exorcise to assess the extent of off-target effects in 55 commercially available CRISPR screen libraries (**Figure 2A**) and found that all inspected libraries contain guides with off-target effects.

**Figure 2.**
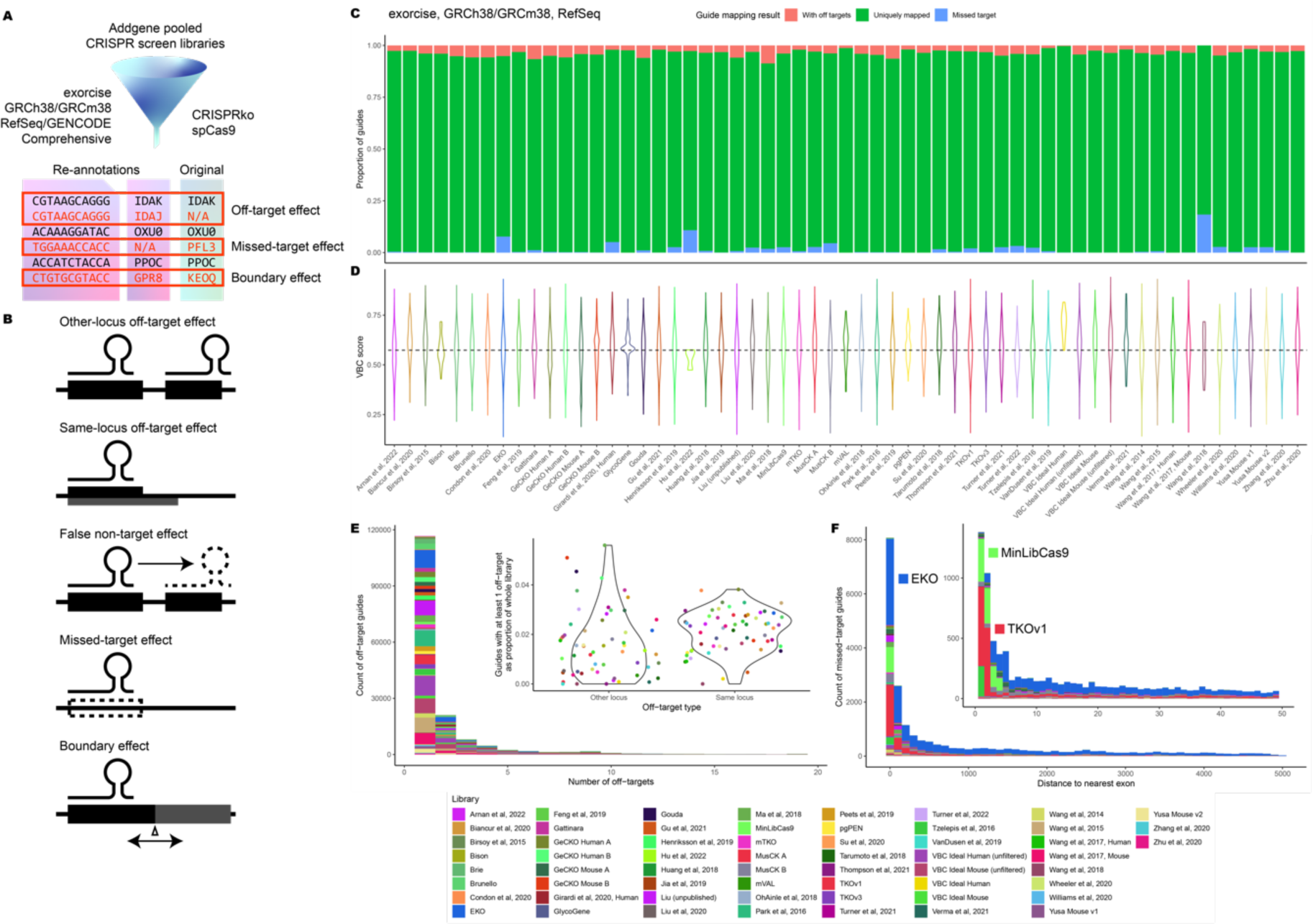
Assessment of Addgene pooled CRISPR-spCas9 libraries for human and mouse. A) Library guides were reannotated with Exorcise using genome assembly GRCh38 (human) or GRCm38 (mouse). Reannotations were compared with original annotations to identify off-target effects, missed-target effects, and boundary effects. B) Mis-annotations identified by Exorcise. Other-locus off-target effects are caused by multiple targeting of guides to multiple loci. Same-locus off-target effects are caused by multiple knockouts by a guide due to overlapping features at a single locus. False non-targeting effects are missing annotations of a valid guide target. Missed-target effects are caused by a library guide being mis-annotated as targeting. Boundary effects are caused by mis-annotations with adjacent exons or genes. C) Performance of Addgene libraries. Library guides were analysed with Exorcise with RefSeq exons. Bars indicate the proportion of guides which have any off-target effects (peach), any missed-target effects (cyan), or only on-targets (green). D) Distribution of VBC scores among Addgene libraries. Dashed line indicates the median VBC score across all guides in all libraries. E) Off-target effects of library guides in Addgene libraries. Distribution of number of off-target effects per guide by library. Inset: distribution of other-locus and same-locus off-target effects by library. F) Missed-target effects in Addgene libraries. Distance between Cas9 cut site, if any, and the nearest exon in linear distance in nucleotides. Inset: zoom plot between 0 and 50 nucleotides.

We defined off-target guides as those guides which target exons of more than one gene with perfect complementarity (**Figure 2B**). We found that off-target effects account for up to 7.4% of library guides within RefSeq exons (**Figure 2C**, **Supplementary Table S1**), rising to 12.9% after Exorcise with GENCODE Comprehensive (**Supplementary Figure S1A**, **Supplementary Table S33**). Since both overlapping gene features (such as gene bodies with read-through transcripts and antisense RNAs) and gene duplication events would both be counted as off-target effects, we next decomposed off-target effects into “other-locus” and “same-locus” effects: the former resulting from Cas9 recruitment to multiple exonic loci and the latter resulting from Cas9 recruitment to a single exonic locus. Other-locus off-target effects did not exceed 5.1% of total library guides in either RefSeq (**Figure 2E**, **Supplementary Tables S2 and S3**) or GENCODE (**Supplementary Figure S1B**, **Supplementary Tables S34 and S35**) annotations; but same-locus off-target effects account for up to 2.2% and 9.3% of library guides for RefSeq and GENCODE, respectively. This is due to the permissive nature of GENCODE Comprehensive annotations, which include lower-confidence transcripts that are not included in RefSeq, resulting in more features at loci but not more loci with features. In general, off-target effects are expected due to the repetitive nature of sequences that underwent gene duplication.

Since guide reannotation by Exorcise is agnostic to prior annotations and is determined by genome alignment and exome specification alone, it is meaningless to refer to an on-target gene when considering a guide’s off-target effects – all targets would be equally valid, barring guide efficiency differences^5–7^. For this reason, off-target guides in CRISPR libraries for which only one of all the possible valid targets is annotated consequently have missing annotations for all the other valid targets. We term these missing annotations “false non-targeting effects”.

Therefore, we tolerate the design of guides which target more than one gene provided that the analysis is aware of all targets. Omitting a guide simply because it has off-target effects is not a valid strategy given the non-random nature of genome evolution and the restrictive nature of CRISPR guide design, which requires consideration of a PAM. Including a guide that has off-target effects but omitting annotations with its off-targets introduces false non-targeting effects in which a guide exists that targets a gene, but its signal is ignored. Exorcise reannotates guides with all the targets that it detects, thereby eliminating false non-targeting effects.

### Missed-target effects are more prevalent in libraries using permissive design strategies

Next, we asked whether any targeting guides in the 55 commercially available libraries miss their targets. We defined missed-target guides as those guides that do not target any exons. Exorcise with RefSeq revealed missed-target effects account for up to 16.1% of library guides, falling to 9.6% after Exorcise with GENCODE Comprehensive. This difference is expected because of the additional lower-confidence annotations available in GENCODE but not RefSeq. We found that this fraction falls most sharply in libraries using permissive design strategies. For instance, the EKO library was designed on putative protein-coding regions in AceView and GENCODE, and so Exorcise of EKO against GENCODE recovered guides that are missed targets in RefSeq. However, even after Exorcise with GENCODE, EKO still contains 7.8% missed-target guides.

To test whether missed-target guides were due to mis-design, we measured the distance between the computed cut site for each missed-target guide target, if any, and the nearest exon. We found a tendency for missed targets to have cut sites within 100 nucleotides from the nearest exon, with marked enrichments flush against and one nucleotide away from an exon boundary (**Figure 2F**, **Supplementary Table S4**). This was most strongly observed in the MinLibCas9 and TKOv1 libraries and the observation was retained after Exorcise with either RefSeq or GENCODE (**Supplementary Figure S1C**, **Supplementary Table S36**). The presence of missed targets within this interval indicates either inconsistent exon boundaries between references, or a design decision to accept cut sites outside but adjacent to an exon boundary as opposed to within it. Cut sites not explicitly within the bounds of an exon enable the possibility that indels acquired by repair at the cut site might leave the exon boundary, and therefore the sequence, the transcript, and the protein product, intact, thereby violating the assumption that successful CRISPR targeting results in a genetic knockout.

The presence of missed-target effects violates the assumption that introduction of targeting guides causes a change in the coding sequence of a gene. Missed-target effects can occur due to design of guides with excessively permissive reference sets or by accepting cut sites at exon boundary positions where the repair outcome is unclear. In both cases, targeting may be successful but would not translate to a detectable phenotype. Since subject cell lines, especially those with genomic instabilities such as cancer cell lines^8^, are unlikely to reflect every CRISPR target identified in permissive reference exomes with low-confidence transcripts, we recommend that CRISPR targets be verified in the cell line to be investigated. Exorcise helps with this by re-annotating missed-target guide RNAs given a user-defined cell line genome and exome, thus assuring the assumption that guides in the library possess valid CRISPR targets in the cell line under investigation.

### De novo library design with re-annotation retains favourable on-target efficacy

Re-annotation by Exorcise identifies missed-target effects that are removed from the library in the reannotation. When using Exorcise for library design, missed-target guides should be replaced with on-target guides to ensure constant numbers of guides per gene. We therefore asked whether a library designed with Exorcise would retain on-target efficacy. We obtained and re-annotated the top 20 guides per gene by Vienna Bioactivity CRISPR (VBC) score, an on-target metric for CRISPR guide design, in human GRCh38 and mouse GRCm38 genomes, separately, using RefSeq exomes. Exorcise revealed very few missed-target guides due to VBC itself being designed on RefSeq exons; we removed them as well as all other-locus off-target guides. From the remaining guides, we accepted the top six guides per gene by VBC score into new libraries, which we designate “VBC Ideal Human” and “VBC Ideal Mouse”, respectively (**Supplementary Tables S5 and S6**). These libraries had distributions of VBC scores and off-target fractions comparable with the other libraries we assessed (**Figure 2D**). Because we explicitly removed other-locus off-target guides in the design of the libraries, this fraction decreased after moving from 20 guides per gene to six. Subsequent Exorcise of the final libraries with GENCODE Comprehensive exomes retained freedom from missed-target guides, while off-target fractions increased. However, this is expected due to the GENCODE reference being more permissive. Taken together, we show that Exorcise is an attractive method for validating CRISPR guide targets for library design due to its ability to identify and correct off-target and missed-target effects. Combining Exorcise with library design yields balanced libraries with a uniform number of guides per gene while validating CRISPR guide targets.

### Simulations of mis-annotation reveal impacts on CRISPR screens

To appraise the effects of common mis-annotations in CRISPR guides on screens, we generated a synthetic chemo-genetic dataset with prescribed gene-drug interaction values (**Figure 3A**, **Supplementary Table S7**). We assigned guides to genes with either the ground truth or one of four mis-annotated exome schemes (**Supplementary Tables S8 and S9**) and challenged each scheme to capture the prescribed interactions from simulated CRISPR data (**Supplementary Table S10**). The ground truth scheme mapped guides to their intended gene uniquely, prescribing three targeting guides per gene. A “false non-targeting” scheme randomly switched targeting guides as non-targeting, resulting in between one and three guides being assigned per gene. A “missed targets” scheme assigned the same three correct guides per gene plus up to seven additional non-targeting guides per gene. A “boundary” scheme randomly shifted the exon boundaries into adjacent gene bodies such that each gene may be assigned up to seven guides, of which up to four may be off-target guides. Finally, a “random” scheme was designed where mappings were randomly created between guides and genes, even if this resulted in discontinuous gene bodies.

**Figure 3.**
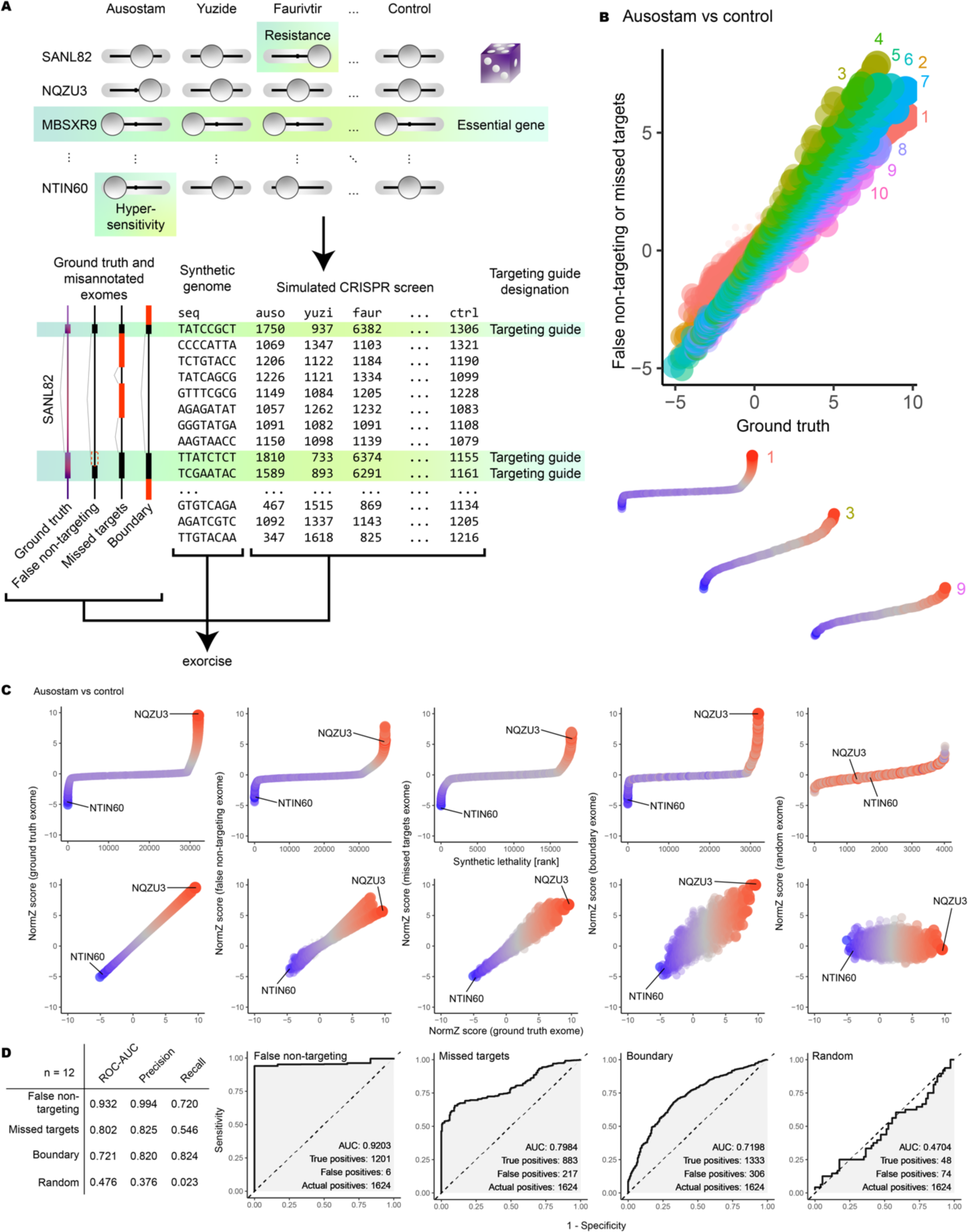
Simulated CRISPR screen on synthetic data, DrugZ analysis. A) Upper: chemo-genetic interaction values were randomised for each gene-drug pair across 4000 genes and 12 drugs. A control drug was defined with chemo-genetic interaction value 1. Essential genes were modelled with chemo-genetic interaction value 0.1 with all drugs and the control drug. Lower: A synthetic genome was constructed with ten guides per gene, three of which targeting exons. A ground truth exome was constructed that reflected the exon structure of each gene correctly. Mis-annotated exomes were constructed: a false non-targeting exome excluded targeting guides from exons; a missed-targets exome included non-targeting guides into exons; a boundary effects exome included guides from adjacent genes into exons. Red regions in the mis-annotated exomes indicate differences from the ground truth. CRISPR read counts were simulated according to chemo-genetic interactions and targeting guide designations. Simulated read counts were annotated with Exorcise using the synthetic genome and the ground truth or mis-annotated exomes. B) Upper: Bundle analysis. DrugZ normZ scores for simulated gene knockouts in the simulated drug ausostam versus control. Each point is one gene. Small points indicate essential genes. Shown are genes annotated with the ground-truth exome versus those annotated with the false non-targeting exome (labels 1 and 2), the ground truth exome (label 3), or missed-targets exome (labels 4-10). Labels indicate number of guides per gene: 1 and 2 indicate missing targeting guides (false non-targeting); 4-10 indicate additional non-targeting guides (missed targets). Lower: example rank plots for genes represented at 1, 3, and 9 guides per gene, plotted on identical scales. C) Simulated CRISPR screen as in (B). Upper: rank plots for each exome annotation. Lower: biplot of normZ scores of genes with each exome annotation versus ground truth. NQZU3 (ground-truth resistance signal) and NTIN60 (ground-truth hypersensitivity signal) are shown in all plots. Points are coloured by chemo-genetic interaction value: red, resistance; blue, hypersensitivity. Small points indicate essential genes. Ausostam shown; representative results across 12 independent simulations. D) Left, summary statistics of the receiver-operator characteristic (ROC) curve analysis. Right, ROC curves showing the performance of mis-annotated schemes to recover the discoveries made by the ground truth scheme. AUC, area under the ROC curve. Ausostam shown; representative results across 12 independent simulations.

As expected, the ground truth scheme captured the correct chemo-genetic interactions after DrugZ (**Figure 3C**, **Supplementary Figures S2 & S3**, **Supplementary Tables S15–S24**) and MAGeCK (**Supplementary Figures S4–6**, **Supplementary Tables S37–S46**) analysis of the dataset. The boundary scheme could only capture the strongest interactions and discovery of weaker interactions was impaired. The missed targets and false non-targeting schemes recovered chemo-genetic interactions best when the number of guides per gene was similar to that in the ground truth scheme; that is, three. Mis-annotation by introduction of non-targeting guides into exons or removal of targeting guides from exons impaired recovery of chemo-genetic interactions. Finally, as expected, the random scheme did not successfully capture chemo-genetic interactions.

In missed targets and false non-targeting schemes, we found that super-numerary and sub-numerary guides per gene affected discovery strength but preserved discovery direction (that is, conferring drug hypersensitivity or resistance) (**Figure 3B**, **Supplementary Tables S11–S14**). This was observed on plots of ground truth versus mis-annotation normZ scores as a “bundling” effect (**Figure 3C**) created by loci of normZ scores falling on straight lines with different gradients. Individual straight lines within a bundle represented genes with a distinct number of guides per gene. Bundle line gradient, and therefore discovery strength, increased as guides per gene approached the ground truth value of three. Deviations from three guides per gene due to addition of non-targeting guides or removal of targeting guides attenuated discovery strength but largely preserved direction and order.

We next quantified discovery strength by considering receiver-operator characteristic (ROC) curves for mis-annotated schemes to act as classifiers for the ground truth. We defined actual positives by using the ground truth scheme and challenged each mis-annotated scheme to recover the actual positives and exclude false positives (**Figure 3D**). The false non-targeting scheme performed the best, with an average area under the ROC curve (ROC-AUC) of 0.932 over 12 independent simulations. It recovered actual positives almost to the exclusion of false positives (precision = 0.994) but did not recover all of the actual positives (recall = 0.720). The missed targets scheme performed worse, recovering only the very strongest actual positives before supernumerary guide dilution impaired exclusion of false positives. The boundary scheme performed the worst (ROC-AUC = 0.721) apart from random control, although it did recover the most actual positives (recall = 0.824). Taken together, we found that mis-annotations that introduce additional guides per gene – that is, missed targets and boundary effects – represent the largest penalty to discovery strength. Mis-annotation by omission of targeting guides (false non-targeting) reduces the number of discoveries but does not impair discovery precision, and this demonstrates why libraries designed with few guides per gene, such as Gattinara, still perform well.

Taken together, as demonstrated by simulations, all common mis-annotations have an adverse effect on discovery. Mis-annotations that introduce incorrect guides have a larger adverse effect than mis-annotations that remove correct guides. We therefore recommend that CRISPR libraries be validated for the cell line under investigation to ensure that valid CRISPR targets in the cell line genome are considered appropriately to avoid false non-targeting, missed target, and boundary effects. Exorcise validates CRISPR library guides by alignment to a user-defined genome.

### Reannotation of DepMap cancer cell line CRISPR screens

Next, we demonstrated the applicability of Exorcise on generating personalised reannotations of library guides on cancer cell lines in the Cancer Dependency Map (DepMap)^9^. Cancer cells undergo genomic rearrangement and instability, so it is inadequate to assume that all CRISPR targets designed on standard genome assemblies such as GRCh38 are valid in a cancer genome. We addressed this issue by deducing cancer cell line exomes from RNA-seq data and re-annotating based on transcript abundance (**Figure 4A**). We assumed that transcripts expressed to at least one transcript per million reads (TPM) were present in the exome, and if so, then we included the associated RefSeq exons for those transcripts into the exome. We compared this TPM-based strategy with a parallel strategy using all RefSeq exons regardless of transcript abundance and compared both with the published CRISPR dependency scores on DepMap.

**Figure 4.**
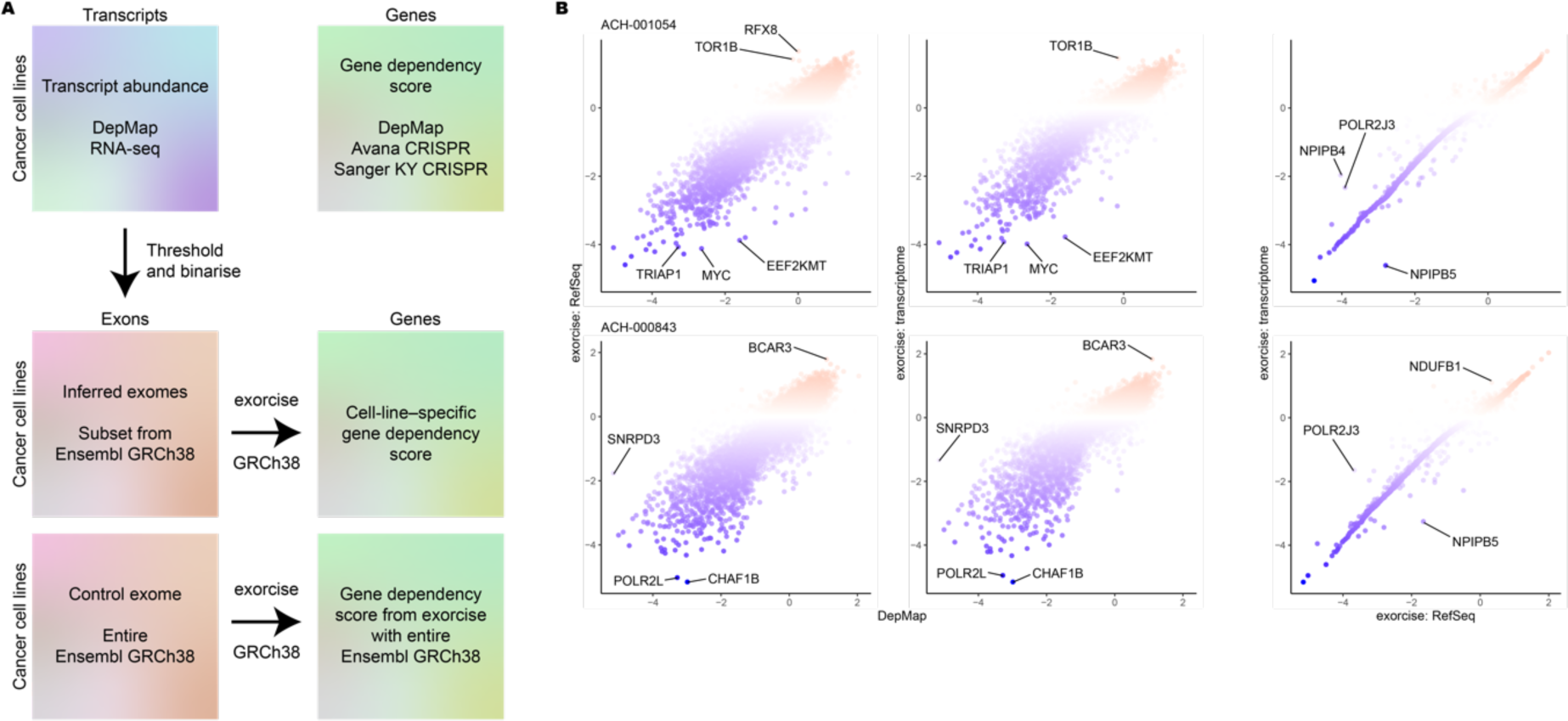
Reannotation of DepMap CRISPR screens with transcriptomes. A) Schematic of exome inference from transcriptomic data and reannotation with Exorcise. B) Representative example of re-annotated DepMap CRISPR screens. Values indicate normalised gene dependency scores (DepMap) or normalised normZ scores (Exorcise). Differential genes are highlighted. Points are coloured by normalised gene dependency score on the y-axis.

Both TPM-based and RefSeq exome strategies agreed strongly with DepMap dependency scores, indicating retention of discoveries regardless of strategy (**Figure 4B**, **Supplementary Tables S25 and S26**). Between the two strategies, we saw strong retention of discoveries at the extreme tails but differences in discoveries with intermediate normZ scores. Uplift in the magnitude of intermediate normZ scores could indicate correction of missed target effect guides that were absent in the cancer genome as evidenced by low TPM expression.

Taken together, we demonstrate concordance between gene dependency scores computed by Exorcise and published on DepMap. We further demonstrate that exome estimation for Exorcise re-annotation is possible from transcriptomics where genomics is not available. We posit that Exorcise corrects missed target effects by excluding guides in the library when evidence from TPM expression suggests that the target is not expressed and therefore is not in the exome. However, since lack of transcript expression does not directly indicate whether the target exists in the genome, whole genome sequencing is required to validate this assumption.

### Reannotation of published DDR CRISPR screens identifies improved signal in intermediate hits

Finally, we explored whether existing DNA damage response (DDR) CRISPR screens would benefit from reanalysis with a library reannotated by Exorcise. We subjected all of the CRISPR screens covered in the DNA damage response CRISPR screen portal (DDRcs)^4^ to Exorcise with RefSeq GRCh38 and compared DrugZ normZ scores with and without Exorcise. Across all screens, we found 105 genes that exhibited an absolute normZ improvement of at least 3 (**Figure 5A**, **Supplementary Table S27**). Among these 105 genes were multiple members of gene families, for example, *NBPF*, *NPIPA*, *PRR20*, *TBC1D3*, and *USP17L*, whose appearance is not surprising given the extent of sequence identity among family members, leading to numerous false non-targeting mis-annotations and improved number of guides per gene after reannotation. However, we also observed benefit in genes that presented in the absence of family members, such as Aicda, *EIF3C*, Polr3k, and *TAF9*.

**Figure 5.**
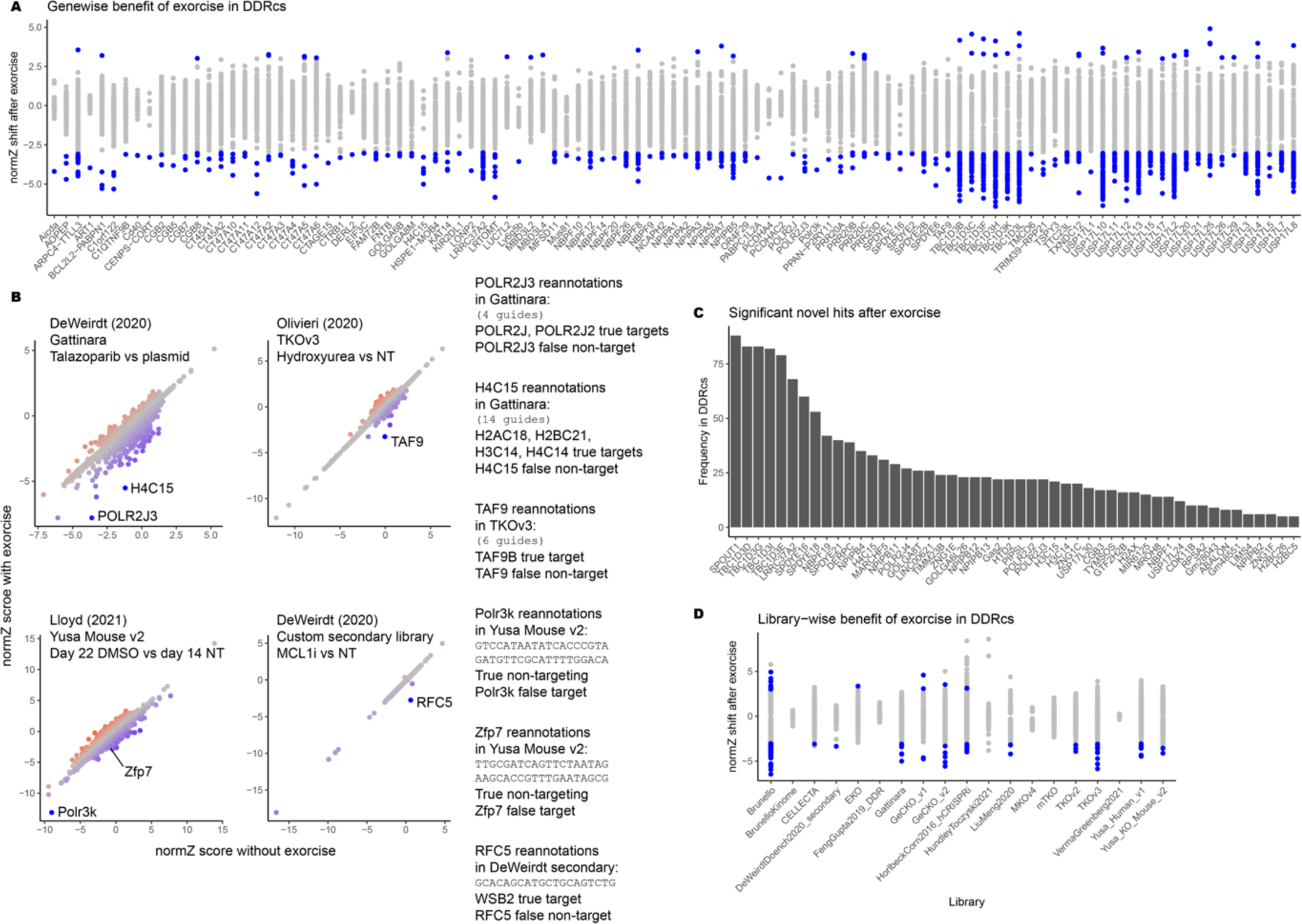
Reannotation of DDRcs with select examples. A) DrugZ normZ score shift (Exorcise normZ – original normZ) by gene. Each point indicates one experiment in the DDRcs. Shown only are genes in which at least one experiment has a normZ shift greater than 3 or less than −3. Blue points indicate experiments in which that gene had a normZ shift greater than 3 or less than −3. B) Bi-plots of select reannotated experiments in the DDRcs. Clockwise from top-left: talazoparib from experiment start, Gattinara library, DeWeirdt (2020)^15^; hydroxyurea (acute) at matched timepoints, TKOv3 library, Olivieri (2020)^11^; MCL1 inhibitor S63845 at matched timepoints, DeWeirdt secondary library, DeWeirdt (2020)^15^; DMSO day 22 versus no treatment day 14, Yusa Mouse library, Lloyd (2021)^10^. Blue points indicate more negative normZ shift (decreased with Exorcise); red points indicate more positive normZ shift (increased with Exorcise). Text on right shows per-guide reannotation details of highlighted differential hits. C) Genes that became significant (DrugZ FDR ≤ 0.05) with Exorcise but not without. Pseudogenes, readthrough transcripts, antisense transcripts, and uncharacterised genes are excluded. D) DrugZ normZ score shift as in (A) but by library. Blue points indicate gene/experiment pairs using that library with a normZ shift greater than 3 or less than –3 in the same direction as without Exorcise.

Polr3k, for instance, showed improved negative normZ score in the untreated case after Exorcise in the Yusa Mouse V2 library due to two of its guides being reannotated from targeting to non-targeting (**Figure 5B**, **Supplementary Tables S28-S31**). Our original analysis had identified Polr3k as an essential gene where depletion resulted in a fitness defect^10^. Analysis after Exorcise reannotation suggested a stronger fitness defect.

*TAF9* in the TKOv3 library was reannotated to include additional guides for *TAF9B* that also target *TAF9* (a false non-targeting mis-annotation) to improve the discovery of *TAF9* hits. In a screen for resistance and sensitivity to acute hydroxyurea treatment, the authors identified mostly non-DDR genes as hits in their original analysis^11^, but not the accessory transcription factor gene *TAF9*. After Exorcise, analysis revealed *TAF9* depletion as hypersensitising to acute hydroxyurea treatment, consistent with hydroxyurea’s role in inducing transcriptional changes related to activation of the DDR^12–14^.

In Gattinara, guides targeting *POLR2J* and *POLR2J2* also target *POLR2J3*. Exorcise corrected these false non-targeting mis-annotations so that they also target *POLR2J3*. Furthermore, 14 guides targeting *H2AC18*, *H2BC21*, *H3C14*, and *H4C14* were reannotated to correct false non-targeting mis-annotations for *H4C15*. In a screen for sensitisers and resistance to talazoparib, using the Gattinara library^15^, both *POLR2J3* and *H4C15* exhibited more negative normZ scores after Exorcise. Although neither of these genes were hits in the original screen, their relevance in the reannotation should be considered with care due to the large increase in guides representing each gene compared to the two guides per gene design of Gattinara.

By the same authors, another screen inspecting the effect of CRISPR knockouts in MCL1 inhibited cells with a custom secondary library in which intermediate sensitising hits in Meljuso and OVCAR8 cells included genes involved in ribosome biogenesis, cell cycle checkpoints, or ubiquitylation^15^. Exorcise reannotation revealed a new intermediate hit, *RFC5*, for which one guide targeting *WSB2* in the DeWeirdt secondary library was also a false non-target for *RFC5*. The appearance of this gene resulting in moderate sensitisation to MCL1 inhibition is consistent with the other hits originally identified.

Across all reannotated experiments in the DDRcs, we were able to identify genes that in multiple experiments became significant hits only after Exorcise (**Figure 5C**, **Supplementary Table S32**). Among these genes were multiple members of the same family (for example, *TBC1D*, *SPDYE*, and *NPIPB* families) for which reannotation enabled correction of false non-targeting errors among family members, thereby increasing the number of guides representing the same gene in the analysis. The appearance of genes in the absence of other family members (for example, *SPOUT1*, *DERPC*, and *TIMM23B*) indicated a benefit of Exorcise not related to correction of false non-targeting errors between family members.

We also investigated whether some libraries benefitted more than others after Exorcise by inspecting the distribution of normZ shifts across all experiments in the DDRcs with library (**Figure 5D**, **Supplementary Table S27**). We found that almost all libraries had at least one experiment in which at least one gene had a normZ shift of at least three units in the original direction after Exorcise, indicating universal applicability of the algorithm. We also found a bias towards discovery of hits on the hypersensitising side of the analysis. This was an effect also seen in our simulations, where the strongest hits and their ordering on the hypersensitising side were more sensitive (**Figure 4C**) and less susceptible to missed discovery due to mis-annotation (**Figure 4D**). We believe that this is an artefact of the DrugZ analysis tool selected, as we do not see this bias in analysis of the same simulations using MAGeCK (**Supplementary Figure S4**).

Taken together, our reannotations of published screen data indicate a benefit by Exorcise for the enhanced discovery of intermediate hits. We have demonstrated that those hits may hold relevance in the context of stronger hits, which are retained after Exorcise, and we posit that they should not be ignored. Exorcise is able to reveal these intermediate hits.

## Discussion

CRISPR libraries are designed on standard genome assemblies, but CRISPR screens are performed on diverse cell lines. It is not appropriate to assume that all guides have valid targets in every cell line under investigation, and this is particularly true for cancer cell lines. Exorcise fills the unmet need for cell-line–specific CRISPR guide design without requiring costly bespoke library design or alteration of CRISPR screen protocols. In this work, developed Exorcise to inspect 55 commercially available libraries on Addgene for common mis-annotations relative to the restrictive RefSeq and permissive GENCODE Comprehensive reference sets. Finding that false non-targeting, missed-target, and boundary effects were common mis-annotations, we then simulated a synthetic CRISPR screen and tested mis-annotated exomes for their ability to recover the ground truth. Confirming that super- and sub-numerary guide annotations contribute to recovery strength but not direction, we then reannotated DepMap CRISPR screen guides, using the transcriptome as a proxy for the exome, to eliminate mis-annotations while preserving the original discoveries and demonstrate universal benefit of Exorcise across libraries and screens in the DDRcs. We therefore present Exorcise as a powerful tool to validate CRISPR library guides for the cell line under investigation by alignment to a user-defined genome and annotation with a collection of exons known or deduced to be utilised by the cell. Exorcise is agnostic to all previous annotations, if any, and relies solely on the guide sequence to produce an alignment and re-annotation. If provided with an *a priori* set of gene annotations, Exorcise additionally produces mappings between original and re-annotations, enabling alignment-based harmonisation of gene symbols conforming to the user-defined exome.

Our evaluation of Addgene libraries revealed that all libraries have off-target effects and libraries designed using more permissive reference sets had missed-target effects. Due to repetitive sequences in the genome, off-target effects can be tolerated, but those guides should be annotated with all the targets, and omission of this leads to false non-targeting effects. Since permissive reference sets such as GENCODE Comprehensive include evidence from low-confidence transcripts, targets designed on permissive references are more likely to violate the assumption that targets in the library exist in the cell line under investigation. For future CRISPR library guide design, we recommend using restrictive reference sets such as RefSeq to ensure the widest universality of valid targets across cell lines even if Exorcise is not to be performed.

We constructed VBC Ideal Human and VBC Ideal Mouse libraries using Exorcise to identify and remove other-locus off-target effects and account for remaining false non-targeting effects relative to RefSeq exons. Using Exorcise at the library design stage enables the investigator to validate CRISPR guide targets in their desired cell line and verify their validity across cell lines. Exorcise reannotation introduces changes to the numbers of guides per gene represented in the library, which we show through simulations results in differential discovery power across genes. Therefore, using Exorcise at the library design stage is ideal to simultaneously validate guides while ensuring that library gene representation remains balanced.

In simulations, we tested the ability of libraries mis-annotated with missed-target, false non-targeting, and boundary effects to recover ground-truth simulated chemo-genetic interactions and found that reducing the number of targeting guides from exons or introducing non-targeting guides into exons each attenuated recovery. These simulations show that mis-annotations observed in commercially available CRISPR libraries could potentially hinder discoveries with intermediate magnitude. Exorcise corrects these mis-annotations, removing the annotations of missed-target guides and permitting multiple annotation of off-target guides where their omission would cause false non-targeting effects. However, our simulations also show a possible effect of non-uniform numbers of guides per gene even when no mis-annotations exist, as would be the case after Exorcise. We found that DrugZ normZ scores for genes re-annotated resulting in fewer guides per gene were consistently smaller than those with more guides per gene. Although this might obscure discoveries of genes with lower representation by the coincidence that reannotation reduced their representation, we believe that this approach is statistically consistent as one’s confidence in the evidence for a discovery increases as the number of observations – that is, guides per gene –, increases.

Our reannotation of DepMap screens showed that *de novo* genome assembly and determination of exome coordinates on that assembly are not necessary for execution of Exorcise for bespoke cell line genomes. We showed that transcriptome TPM counts provide sufficient evidence to deduce the presence or absence of exons in the cell line under investigation and that this information can refine a pre-existing exome on a pre-existing genome assembly. Although this approach misses novel genes in the cell line genome under investigation, it is unlikely that such novel genes would be covered by commercially available CRISPR libraries.

Our reannotation of DDRcs screens agreed with DepMap reannotations and simulations, in that Exorcise can reveal and strengthen intermediate screen hits where the evidence from guide alignment to the genome exists. We found that missed-target and false non-targeting mis-annotations can be reannotated to reflect all valid targets for that guide, increasing the number of guides for that gene and providing better evidence for its chemo-genetic effect. We found that reannotation of both effects enables the discovery of hits that were not significant without Exorcise. We observed a bias for new discoveries to be made on the hypersensitivity side of screens and reconcile this as an artefact of the DrugZ analysis method, as we also saw this effect in DrugZ analysis of our simulated CRISPR screen but not in MAGeCK analysis.

Taken together, we show that Exorcise validates CRISPR library guides to the target genome of the user’s choice. The algorithm corrects false non-targeting, missed-target, and boundary effects and we establish that these are among commonly occurring mis-annotations in commercially available CRISPR screen libraries. Through reannotation, Exorcise enables improved evidence for discovery in the intermediate range while preserving original discoveries. Exorcise is compatible with arbitrary user-defined genome assemblies and exome boundary specifications to the same assembly coordinates, and transcriptome data are sufficient to refine a standard exome for this purpose. It has not escaped our notice that Exorcise can be used to annotate arbitrary DNA sequences by alignment to the genome and therefore has wider potential uses beyond CRISPR screens.

## Materials and methods

### Exorcise algorithm

Briefly, guides are aligned to the supplied genome, a cutting interval is determined, and exon hits are recorded as overlaps between cutting intervals and the exome. Guides inherit the annotations from the exome at the positions of those overlaps.

Guides are appended at the with the *Streptococcus pyogenes* (sp) Cas9 protospacer adjacent motif (PAM) sequence “NGG” at the 3’ terminal end of the guide. BLAST-like alignment tool (BLAT)^16^ is run on the list of guides against a user-supplied genome with the parameters “-stepSize=4 -tileSize=10 -fine -repMatch=2000000 - minScore=20 -minIdentity=100” to identify the genomic coordinates creating perfect alignments between guides plus PAM and the genome. For each perfect match, genomic coordinates of cutting intervals are determined, defined as a zero-width interval located three and four nucleotides 5’ from the start of the PAM. Where cutting intervals overlap with exons in the user-supplied exome, the guide producing the overlap is annotated with the gene symbol of that exon as specified by the user-defined exome. Guides are permitted to be annotated with more than one gene symbol. Guides not annotated with any gene symbols are instead annotated as non-targeting guides. The collection of reannotated guides is output as the guide-level re-annotation of the library.

Additionally, a gene-level mapping between the library author’s original gene symbol and reannotated symbols is created if the library author’s original guide annotations are supplied. If so, then for each unique original annotation, all reannotations of guides originally annotated with that original annotation (“candidates”) are inspected to create a mapping between the original annotation and the reannotation. If there are no candidates, then no mapping is made. If there is exactly one candidate, then it is accepted. If there are two or more candidates, then the candidate that occurs most frequently is accepted; in the case of a tie, then a candidate is selected according to the following hierarchy (in descending order of preference): same as the original symbol, protein-coding gene, non-coding RNA gene, pseudogene, any other gene or genomic feature. For each original annotation, all guides with that annotation are reannotated with the accepted candidate. The collection of mappings between original annotations, accepted re-annotations, and relevant guides is output as the gene-level reannotation of the library.

In all analyses using Exorcise, version 0.52 was used (see Data and code availability).

### Evaluation of CRISPR libraries

The Addgene pooled CRISPR libraries catalogue (https://www.addgene.org/pooled-library/#crispr, accessed 12-Oct-2022) was searched for presently or previously commercially available CRISPR screen libraries designed for use with spCas9; only libraries targeting human or mouse genes and using CRISPR knockout (CRISPRko) biology were included. Libraries designed for CRISPR activation, CRIPSR inhibition, or base editing were excluded. After filtering, 55 CRISPRko-spCas9 knockout screen libraries were accepted (**Table 1**). Library guide sequences were acquired from Addgene or the respective original literature.

**Table 1.** CRISPR-spCas9 knockout pooled libraries analysed in the study.

| Library | Species | Source |
| --- | --- | --- |
| Arnan et al, 2022 | Human | 23 |
| Biancur et al, 2020 | Mouse | 24 |
| Birsoy et al, 2015 | Human | 25 |
| Bison | Human | 26 |
| Brie | Mouse | 6 |
| Brunello | Human | 6 |
| Condon et al, 2020 | Human | 27 |
| EKO | Human | 28 |
| Feng et al, 2019 | Mouse | 29 |
| Gattinara | Human | 15 |
| GeCKO Human A | Human | 30 |
| GeCKO Human B | Human | 30 |
| GeCKO Mouse A | Mouse | 30 |
| GeCKO Mouse B | Mouse | 30 |
| Girardi et al, 2020, Human | Human | 31 |
| GlycoGene | Human | 32 |
| Gouda | Mouse | 15 |
| Gu et al, 2021 | Mouse | 33 |
| Henriksson et al, 2019 | Mouse | 34 |
| Hu et al, 2022 | Human | 35 |
| Huang et al, 2018 | Human | 36 |
| Jia et al, 2019 | Human | 37 |
| Liu (unpublished) | Human | † |
| Liu et al, 2020 | Mouse | 38 |
| Ma et al, 2018 | Human | 39 |
| MinLibCas9 | Human | 40 |
| mTKO | Mouse | 41 |
| MusCK A | Mouse | 42 |
| MusCK B | Mouse | 42 |
| mVAL | Mouse | 41 |
| OhAinle et al, 2018 | Human | 43,44 |
| Park et al, 2016 | Human | 45 |
| Peets et al, 2019 | Human | 46 |
| pgPEN | Human | 47 |
| Su et al, 2020 | Human | 48 |
| Tarumoto et al, 2018 | Human | 49 |
| Thompson et al, 2021 | Human | 50 |
| TKOv1 | Human | 51 |
| TKOv3 | Human | 52 |
| Turner et al, 2021 | Human | 53 |
| Turner et al, 2022 | Mouse | 54 |
| Tzelepis et al, 2016 | Human | 22 |
| VanDusen et al, 2019 | Mouse | 55 |
| VBC Ideal Human (unfiltered) | Human | This study and <sup>7</sup> |
| VBC Ideal Mouse (unfiltered) | Mouse | This study and <sup>7</sup> |
| VBC Ideal Human | Human | This study and <sup>7</sup> |
| VBC Ideal Mouse | Mouse | This study and <sup>7</sup> |
| Verma et al, 2021 | Human | 56 |
| Wang et al, 2014 | Human | 57 |
| Wang et al, 2015 | Human | 58 |
| Wang et al, 2017, Human | Human | 59 |
| Wang et al, 2017, Mouse | Mouse | 59 |
| Wang et al, 2018 | Mouse | 60 |
| Wheeler et al, 2020 | Human | 61 |
| Williams et al, 2020 | Human | 62 |
| Yusa Mouse v1 | Mouse | 63 |
| Yusa Mouse v2 | Mouse | 22 |
| Zhang et al, 2020 | Human | 64 |
| Zhu et al, 2020 | Mouse | 65 |

† https://www.addgene.org/pooled-library/liu-crispr-knockout/ (accessed 12-Oct-2022)

Guide- and gene-level reannotations were obtained using Exorcise specifying genome assembly GRCh38 for libraries targeting human genes and GRCm38 for libraries targeting mouse genes. Genomes in 2bit format were obtained from the UCSC Genome Browser database^17^ (https://hgdownload.soe.ucsc.edu/downloads.html, accessed 29-Jul-2022). In both cases for human and mouse, both RefSeq^18^ and GENCODE Comprehensive^19^ were specified as the exome, separately. Exomes on the same coordinates as the respective genomes were obtained from the UCSC Genome Browser database^17^ (https://genome.ucsc.edu/cgi-bin/hgTables, accessed 29-Jul-2022).

In the evaluation of Addgene libraries, off-target effect guides were defined as those guides which had more than one unique annotation in the library after Exorcise; same-locus off-target effect guides were defined as those off-target effect guides due to a single alignment; other-locus off-target effect guides were defined as those off-target effect guides due to more than one alignment. Missed-target effect guides were defined as those guides with any original annotation other than non-targeting but a re-annotation of non-targeting. False negative control guides were defined as those guides with an original annotation of non-targeting but any re-annotation other than non-targeting.

### VBC Ideal Human and Mouse library design

GRCh38 and GRCm38 guides were obtained from the VBC resource^7^. The top 20 guides by VBC score per gene symbol with no likely off-targets (number of off-targets OT > 95 equal to zero) were accepted into “VBC Ideal Human (unfiltered)” and “VBC Ideal Mouse (unfiltered)” intermediate libraries for human and mouse genomes, respectively. Exorcise was performed on the respective genomes and RefSeq exons to identify missed-target and off-target effects. No missed-target effects were detected. Guides with other-locus off-target effects were removed and the top six remaining guides per gene by VBC score were accepted into “VBC Ideal Human” and “VBC Ideal Mouse” libraries for human and mouse, respectively.

### In silico simulation

A synthetic genome was constructed containing 4000 simulated genes each with ten CRISPR target sites, of which three were within exons and seven were not within exons. An additional ten CRISPR target sites not within any gene were added. CRISPR guides were designed for all target sites whether within exons (targeting guides) or not (non-targeting guides). Five percent of genes were designated “essential”. Each gene was prescribed random chemo-genetic interaction values with 12 simulated drugs according to Equation 1:

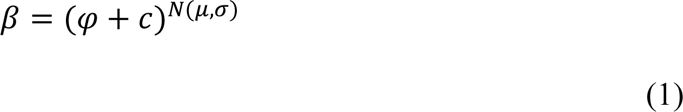

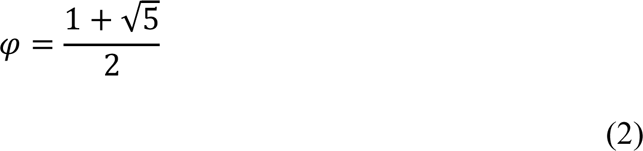

where *β* is the magnitude of the chemo-genetic interaction, *c* is a constant, *φ* is the positive root of the golden ratio (Equation 2), and *N*(*μ*,*σ*) is a random variable with normal distribution centred at mean *μ* and standard deviation *σ*. In the simulation, *c* was set to 0.5, *μ* to 1, and *σ* to 1. These parameters were chosen to ensure a sufficiently wide dynamic range to generate strong and intermediate chemo-genetic interaction values.

Chemo-genetic interaction values smaller than one represent hypersensitivity to the drug when the gene is targeted; values greater than one represent resistance against the drug when the gene is targeted. Genes were also given a control chemo-genetic interaction value representing treatment with a control substance; these values were fixed at one. Chemo-genetic interaction values of essential genes were overwritten with 0.1 regardless of treatment, including the control treatment. Guides inherited the chemo-genetic interaction value of the gene in which they target unless they were non-targeting guides, in which case the chemo-genetic interaction value of those guides was set at one.

CRISPR screen counts were simulated for each guide using guide-level chemo-genetic interaction values according to Equation 3:

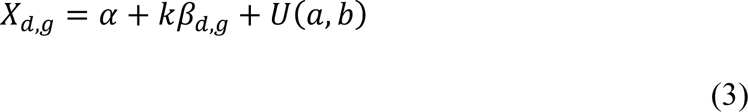

where *X_d,g_* is the read count for guide *g* in drug treatment *d*, *α* is the background read count, *k* is the scaling value representing the number of additional reads per unit chemo-genetic interaction value, *β_d,g_* is the chemo-genetic interaction value between guide *g* and drug treatment *d*, and *U*(*a,b*) is a random variable with uniform distribution and fully closed bounds [*a*, *b*]. In the simulation, *α* was set to 200, *k* to 1000, *a* to -150, and *b* to 150. These parameters were used to ensure that chemo-genetic interaction was the major source of variance and to model random noise in the read counts.

Exomes were created according to one ground-truth scheme, three mis-annotation schemes, and one random scheme. The ground-truth scheme placed exon boundaries such that the three targeting guides per gene were within exons but the seven non-targeting guides per gene were not. That is, the exon boundaries were placed at the correct positions according to the synthetic genome. A “false non-targeting” mis-annotation scheme placed exon boundaries such that up to three targeting guides per gene were randomly selected and place outside of exons instead of within them. A “missed targets” mis-annotation scheme placed exon boundaries such that up to seven non-targeting guides per gene were randomly selected and placed within exons. A “boundary” mis-annotation scheme placed gene boundaries within the gene bodies of up to two neighbouring genes such that up to two targeting guides per gene were placed in exons belonging to the neighbouring gene instead. The random scheme placed exon boundaries randomly across the genome even if it would result in discontinuous gene bodies.

All 40010 simulated CRIPSR guides were aligned to the synthetic genome and annotated with one of the five exomes, each separately, using Exorcise (see Algorithm). Guide-level annotations were transferred to simulated CRISPR read counts. DrugZ differential analysis software^20^ was used to identify simulated genes resulting in hypersensitivity or resistance to each drug treatment compared to control expressed as a normZ score. NormZ scores were compared with the prescribed chemo-genetic interaction values to assess recovery of chemo-genetic interactions in each exome scheme.

Receiver-operator characteristic (ROC) analysis was performed by defining actual positives as genes in the ground truth scheme identified by DrugZ as hypersensitising or resisting with a false discovery rate (FDR) of 0.5 or better. This value was chosen to test the recovery of strong and intermediate signals while excluding low-confidence signals, non-targeting guides, and signals from genes with a chemo-genetic interaction value of close to one. Positives for each of the four mis-annotated schemes were determined by DrugZ using the same FDR cutoff. True positives were defined as those positives which were also actual positives in the same direction. False positives were defined as those positives which were not actual positives or were actual positives but in the opposite direction. ROC curves were determined by considering positives in order of ascending FDR. Area under the ROC curve (AUC) was calculated using the trapezoid method. Precision was calculated as the true positives as a proportion of all positives. Recall was calculated as the true positives as a proportion of actual positives.

### Reannotation of DepMap

CRISPR screen raw read counts from Avana^21^ and Sanger Kosuke Yusa (KY)^22^ libraries and cell line transcriptome transcripts per million (TPM) values were obtained from DepMap^9^ release Public 23Q2 (https://depmap.org/portal/download/all/, accessed 02-Jun-2023). Cell lines used in the CRISPR screens were extracted and those with TPMs were selected. Exomes were deduced from TPMs of selected cell lines by accepting transcripts expressed at 1 TPM or higher. For each selected cell line, CRISPR read counts from that cell line and from the Avana or Sanger KY plasmid, as appropriate, were obtained, guides aligned to GRCh38, and re-annotated with the respective exome using Exorcise. Separately, Exorcise was also performed for each selected cell line with RefSeq genes and non-coding RNAs as the exome. Genes whose knockout modified survival in selected cell lines compared to plasmid were identified using DrugZ. DrugZ normZ scores were compared between transcriptomically-computed exomes, RefSeq exomes, and the published CRISPR gene dependency scores. NormZ scores and gene dependency scores were quantile normalised per screen for display purposes only.

### Reannotation of DDRcs

The DNA Damage Response CRISPR Screen portal (DDRcs)^4^ hosts results from published CRISPR screens interrogating the DDR and provides consistent analysis across all screens including gene nomenclature and differential analysis method. Counts data internal to the DDRcs were obtained, reannotated with Exorcise against RefSeq GRCh38, and differential analysis performed with DrugZ using the same settings as in DDRcs. DrugZ normZ scores were recorded with and without Exorcise alongside the original citation of the screen and the CRISPR library used.

NormZ scores were compared with and without Exorcise for each screen in the DDRcs. *De novo* hits were defined as significant hits in genes that were present with Exorcise but not without, excluding pseudogenes and antisense and readthrough transcripts. Genewise Exorcise benefit was defined as the distribution of differences between normZ scores with and without Exorcise for each gene across all screens. Instances where the sign of the normZ score was different with and without Exorcise were ignored. Library-wise Exorcise benefit was defined as the distribution of differences between normZ scores with and without Exorcise for each library across all genes and screens.

## Supplementary information

**Supplementary Figure S1.**
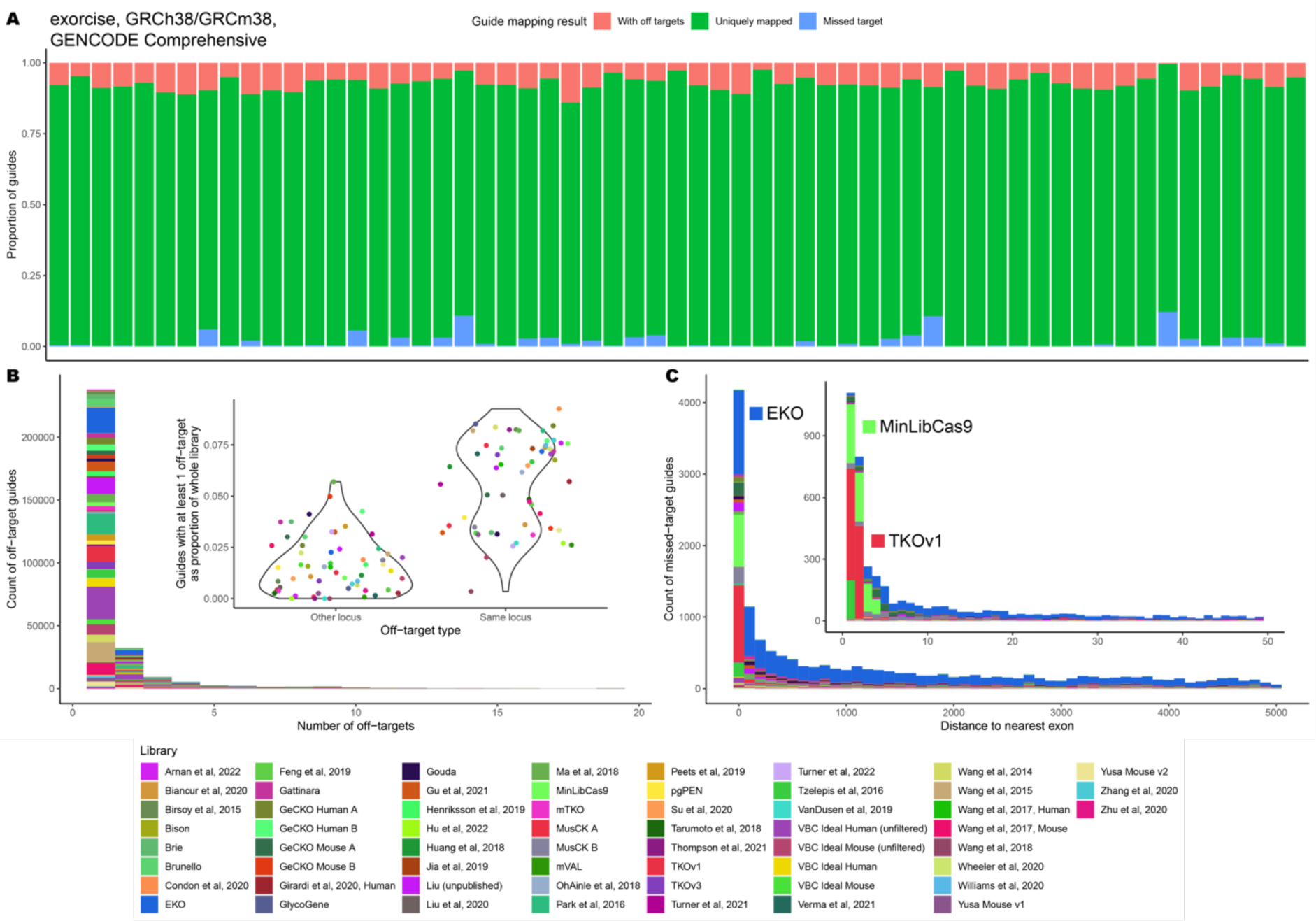
Assessment of Addgene pooled CRISPR-spCas9 libraries for human and mouse using Exorcise with GENCODE Comprehensive. A) As in Figure 2C. B) As in Figure 2E. C) As in Figure 2F.

**Supplementary Figure S2.**
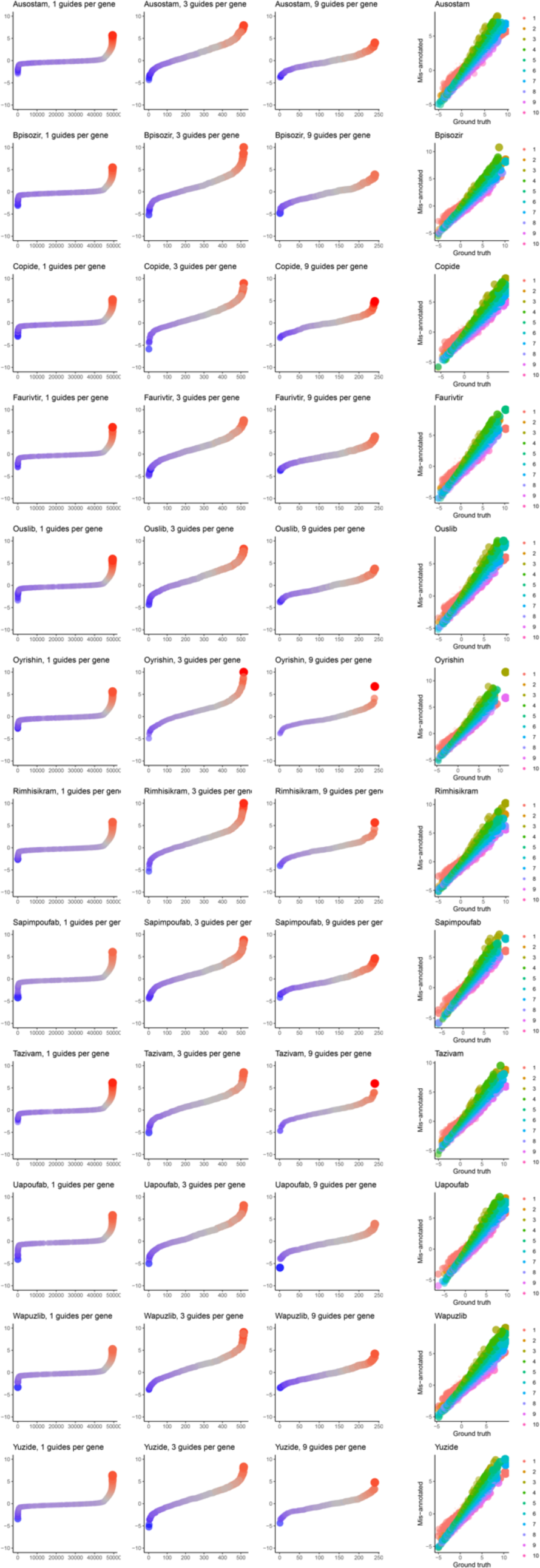
Rank plots and bundle analysis for the simulated mis-annotated CRISPR screen, DrugZ analysis. Rank plots as in Figure 3C upper. Bundle analysis as in Figure 3B upper.

**Supplementary Figure S3.**
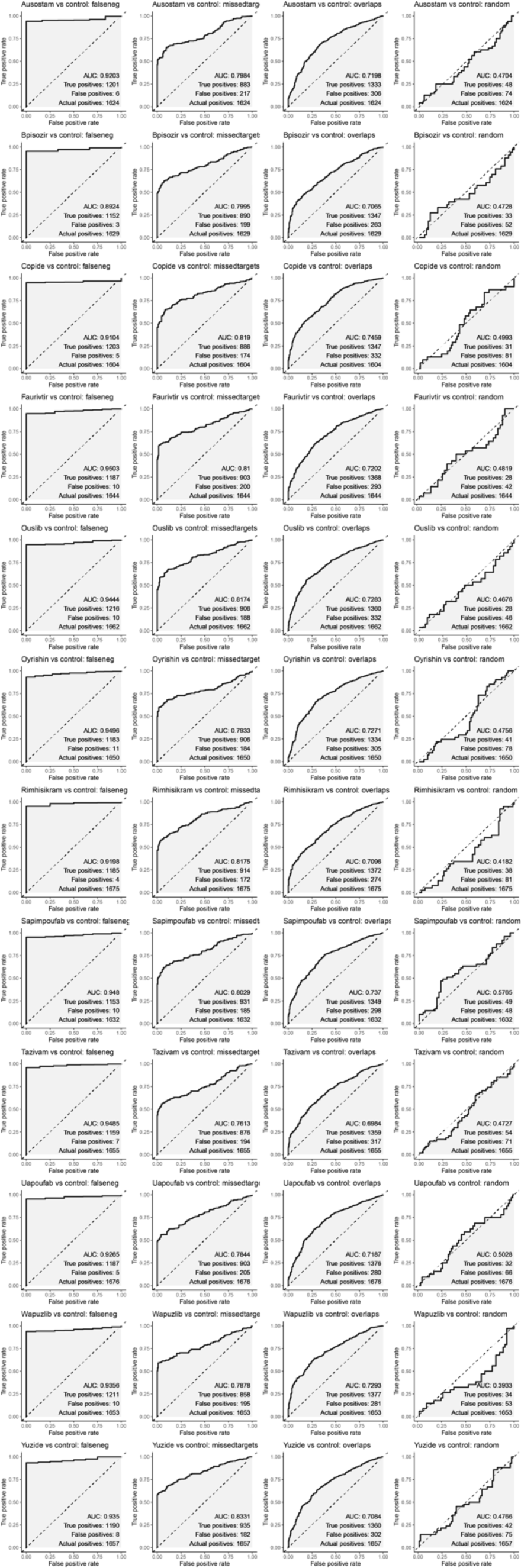
Receiver-operator characteristic (ROC) analysis for the simulated mis-annotated CRISPR screen, DrugZ analysis. As in Figure 3D. Falseneg, false non-targeting mis-annotation scheme; missedtargets, missed targets mis-annotation scheme; overlaps, boundary mis-annotation scheme.

**Supplementary Figure S4.**
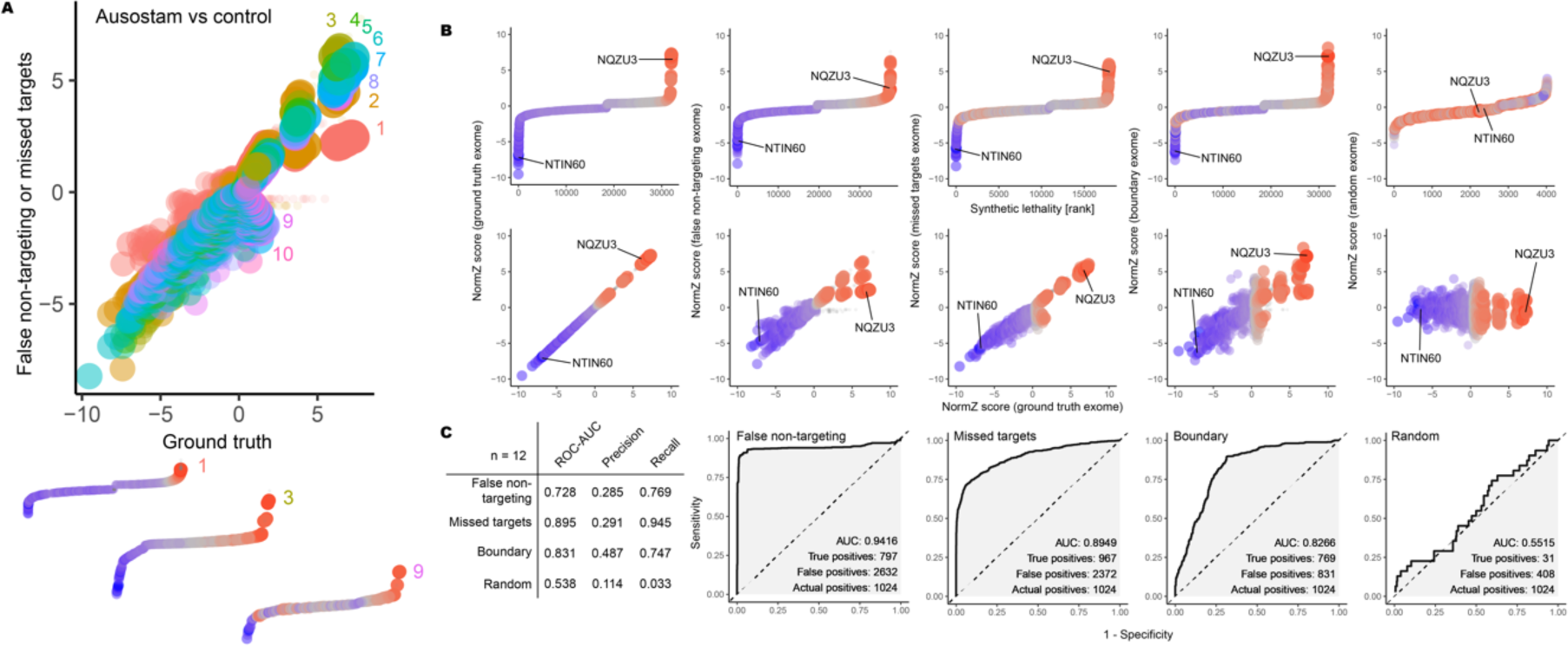
Simulated CRISPR screen on synthetic data, MAGeCK analysis. A) As in Figure 3B. B) As in Figure 3C. C) As in Figure 3D.

**Supplementary Figure S5.**
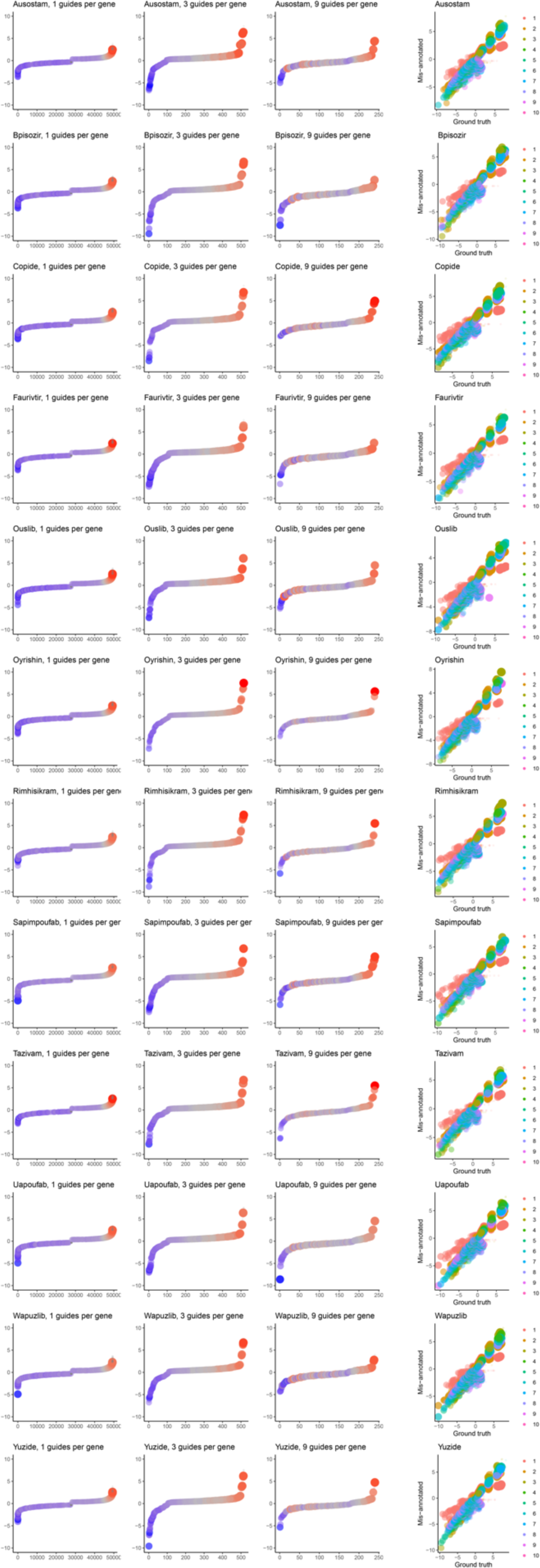
Rank plots and bundle analysis for the simulated mis-annotated CRISPR screen, MAGeCK analysis. Rank plots as in Supplementary Figure S4B upper. Bundle analysis as in Supplementary Figure S4A upper.

**Supplementary Figure S6.**
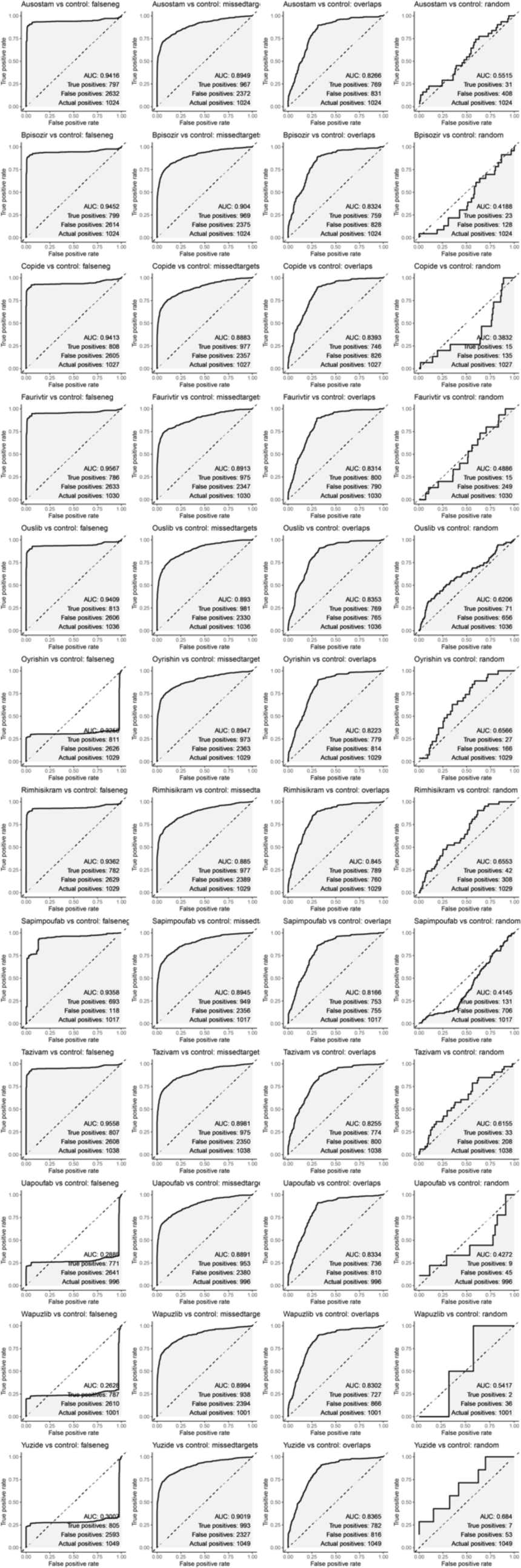
Receiver-operator characteristic (ROC) analysis for the simulated mis-annotated CRISPR screen, MAGeCK analysis. As Supplementary Figure S4C except that the positive classification threshold was 0.9 rather than 0.5.

**Supplementary Table S1.** Source data for Figure 2C.

**Supplementary Table S2.** Source data for Figure 2E.

**Supplementary Table S3.** Source data for Figure 2E inset.

**Supplementary Table S4.** Source data for Figure 2F.

**Supplementary Table S5.** VBC Ideal Human CRISPR-spCas9 knockout guide library after Exorcise with RefSeq.

**Supplementary Table S6.** VBC Ideal Mouse CRISPR-spCas9 knockout guide library after Exorcise with RefSeq.

**Supplementary Table S7.** Simulated chemo-genetic interaction values, related to Figure 3A.

**Supplementary Table S8.** Synthetic guide sequences, related to Figure 3A.

**Supplementary Table S9.** Ground truth and mis-annotated exomes, related to Figure 3A.

**Supplementary Table S10.** Simulated CRISPR screen counts, related to Figure 3A.

**Supplementary Table S11.** Source data for Figure 3B upper.

**Supplementary Table S12.** Source data for Figure 3B lower, trace 1.

**Supplementary Table S13.** Source data for Figure 3B lower, trace 3.

**Supplementary Table S14.** Source data for Figure 3B lower, trace 9.

**Supplementary Table S15.** Source data for Figure 3C upper, panel 1.

**Supplementary Table S16.** Source data for Figure 3C upper, panel 2.

**Supplementary Table S17**. Source data for Figure 3C upper, panel 3.

**Supplementary Table S18**. Source data for Figure 3C upper, panel 4.

**Supplementary Table S19.** Source data for Figure 3C upper, panel 5.

**Supplementary Table S20**. Source data for Figure 3C lower, panel 1.

**Supplementary Table S21**. Source data for Figure 3C lower, panel 2.

**Supplementary Table S22.** Source data for Figure 3C lower, panel 3.

**Supplementary Table S23.** Source data for Figure 3C lower, panel 4.

**Supplementary Table S24**. Source data for Figure 3C lower, panel 5.

**Supplementary Table S25**. Source data for Figure 4B upper.

**Supplementary Table S26**. Source data for Figure 4B lower.

**Supplementary Table S27**. Source data for Figure 5A and 5D.

**Supplementary Table S28**. Source data for Figure 5B upper left.

**Supplementary Table S29.** Source data for Figure 5B upper right.

**Supplementary Table S30**. Source data for Figure 5B lower left.

**Supplementary Table S31**. Source data for Figure 5B lower right.

**Supplementary Table S32**. Source data for Figure 5C.

**Supplementary Table S33**. Source data for Supplementary Figure S1A.

**Supplementary Table S34**. Source data for Supplementary Figure S1B.

**Supplementary Table S35.** Source data for Supplementary Figure S1B inset.

**Supplementary Table S36**. Source data for Supplementary Figure S1C.

**Supplementary Table S37**. Source data for Supplementary Figure S4B upper, panel 1.

**Supplementary Table S38**. Source data for Supplementary Figure S4B upper, panel 2.

**Supplementary Table S39**. Source data for Supplementary Figure S4B upper, panel 3.

**Supplementary Table S40**. Source data for Supplementary Figure S4B upper, panel 4.

**Supplementary Table S41**. Source data for Supplementary Figure S4B upper, panel 5.

**Supplementary Table S42**. Source data for Supplementary Figure S4B lower, panel 1.

**Supplementary Table S43.** Source data for Supplementary Figure S4B lower, panel 2.

**Supplementary Table S44.** Source data for Supplementary Figure S4B lower, panel 3.

**Supplementary Table S45**. Source data for Supplementary Figure S4B lower, panel 4.

**Supplementary Table S46**. Source data for Supplementary Figure S4B lower, panel 5.

## Data and code availability

Exorcise source code is available at <https://github.com/SimonLammmm/exorcise> under a Creative Commons Zero 1.0 Universal licence. Source data are provided within the manuscript and supplementary materials. Additional source data and analysis computer code are available from the authors on request.

## Author contributions

S.L. conceptualised the study, designed the experiments, developed the software, performed the experiments, and analysed the results and wrote the manuscript. J.C.T. provided technical assistance with software development and reviewed the manuscript. S.P.J. reviewed the manuscript and supervised the study.

## Acknowledgements

We acknowledge Dr Tuan-Anh Tran and Dr Sebastian M. Siegner for helpful discussions. We also acknowledge the use of the Cancer Research UK Cambridge Institute high-performance computing cluster provided by the IT & Scientific Computing core facility and thank its staff for the maintenance of the cluster.

